# Super-Resolved Single Small Extracellular Vesicle Assay enabled by a Plasmonic Nanohole Array

**DOI:** 10.64898/2026.06.12.731868

**Authors:** Anthony J. El-Helou, Yiting Liu, Fatemeh Khosravi, Chaohao Chen, Carol H.W. Yan, Mark Lockrey, Juanfang Ruan, Zhaowei Liu, Peter J. Reece, Ying Zhu

**Affiliations:** School of Biomedical Engineering, University of Technology Sydney, NSW 2007, Australia; Electron Microscope Unit, UNSW Sydney, NSW 2052, Australia; Department of Electrical and Computer Engineering, University of California, San Diego, California 92093, USA; School of Physics, UNSW Sydney, NSW 2052, Australia; School of Clinical Medicine, UNSW Sydney, NSW 2052, Australia

## Abstract

The accurate quantification of biological nanoparticles, such as small extracellular vesicles (sEVs), is fundamentally hindered by a resolution-coincidence trade-off in digital assays. While physical confinement can isolate single particles, conventional optical readouts remain diffraction-limited, causing multi-particle occupancy to be miscounted as single events and thereby restricting the analytical dynamic range. Here, we report a nanoplasmonic platform that overcomes this limit by introducing a geometry-defined interface that uniquely unifies nanoscale compartmentalisation and near-field-assisted super-resolution imaging. Utilising a gold plasmonic nanohole array, the strict geometric periodicity of the lattice simultaneously serves as a template for single-vesicle confinement and a deterministic grid that generates an array of localised surface plasmon resonance near-field hotspots. This position-deterministic illumination pattern imposes known geometric priors on the excitation field, shifting high-spatial-frequency information into the detectable bandwidth to achieve sub-100 nm lateral resolution. This dual-purpose geometric determinism enables high-fidelity digital readout of individual vesicles with significantly fewer sub-images than stochastic, speckle-based metasurface structured illumination microscopy approaches. The assay achieves an analytical limit of detection of 143 sEVs/µL, matching the performance of state-of-the-art single-EV counting technologies. It successfully differentiates distinct sEV subpopulations based on surface biomarker expression, establishing a clear pathway for future clinical liquid biopsy applications. By replacing stochastic loading and illumination with geometric design, this work establishes a robust framework for precise vesicle interrogation with broad implications for emerging translational applications and fundamental biology.

## Introduction

Biological nanoparticles, including extracellular vesicles (EVs) and viruses, are emerging as critical tools in biomedicine, with applications ranging from non-invasive diagnostics to precision drug delivery^1^. Small extracellular vesicles (sEVs) represent a class of lipid bilayer-delimited particles released from cells that are typically under 200 nm in diameter^2^. Primarily originating from internal cellular compartments and released via the multivesicular body pathway, sEVs are critical mediators of intercellular communication, mediating the transfer of critical biomolecular cargos that influence both physiology and pathology^3,4^. However, sEVs exhibit significant heterogeneity in their morphology, physiological properties, and biomolecular cargo. Traditional ensemble averaging approaches, such as enzyme-linked immunosorbent assay (ELISA), inherently mask these heterogeneities, leading to misinterpretation of their fundamental roles and practical functions^5^.

Recent advancements in single-EV analysis techniques have begun to unravel sEV heterogeneity^6,7^. Among these methods, digital assays offer a highly sensitive framework by compartmentalising sEVs into discrete partitions, theoretically ensuring a binary occupancy (0 or 1 EVs)^8^. This approach enables discrete counting, which provides superior quantification over ensemble-averaging methods, such as ELISA. Current digital assays achieve analyte compartmentalisation through physical confinement using microwells (e.g., digital ELISA) or droplets (e.g., digital PCR). To minimise coincident occupancy, samples are diluted to follow a Poisson distribution, a “dilute-and-guess’ strategy by statistically discretising the capture events, which restricts dynamic range and throughput^9,10^. While emerging nanoscale confinement strategies have sought to discretise capture events spatially, the subsequent optical readouts remain to rely on diffraction-limited microscopy^11-13^. Consequently, even when analyte particles are physically isolated, stochastic loading at high concentrations can lead to optical coincidence, where multiple particles appear as a single diffraction-limited spot, thereby underestimating counts and compromising analytical accuracy.

Photonic nanohole structures offer a compelling solution by combining spatial discretisation with near-field optical enhancement. While previous studies have used plasmonic nanocavities for Surface-enhanced Raman mapping or Mie-void resonance for colourimetric sensing^14,15^. A platform truly capable of resolving sub-diffraction features within a confined architecture remains unexplored.

Here, we report a super-resolved single-sEV assay that exploits the localised surface plasmon resonance (LSPR) hotspots of plasmonic nanohole arrays (NHAs) to achieve near-field assisted super-resolution imaging with lateral resolution approaching 100 nm. By generating sub-diffraction illumination patterns, the NHA functions as an on-chip structured illumination microscopy (SIM) interface, achieving an approximate three-fold improvement in resolution that surpasses the theoretical two-fold ceiling of conventional SIM. This dual-functional, geometry-defined platform uniquely unifies physical confinement and super-resolution interrogation, enabling precise single-vesicle counting. We demonstrate a limit of detection (LOD) of 143 sEVs/μL and show high analytical selectivity in differentiating sEV subpopulations based on surface biomarker expression. By replacing stochastic sample loading and unpredictable speckle illumination with a lattice-anchored geometric design, this platform establishes a robust framework that successfully overcomes the fundamental resolution-coincidence trade-off in conventional digital assays. This work provides a scalable pathway towards high-fidelity single sEV interrogation with broad implications for future clinical liquid biopsies and fundamental cell biology.

## Results and discussion

### Principle of the NHA-assisted super-resolved single sEV assay

The platform integrates nanoscale compartmentalisation with near-field-assisted SIM to transform stochastic vesicle capture into a geometry-defined, super-resolved readout. The NHA architecture consists of a hexagonal lattice of nanoholes (90 nm diameter; 225 nm pitch) milled through an 80-nm thick gold film deposited on a fused silica substrate (Figure 1A inset). The geometry is precisely engineered to align the plasmonic resonance with the excitation wavelength (Figure S1), generating near-field illumination patterns that contain spatial frequency components beyond the diffraction-limited cutoff^16^. This configuration facilitates efficient frequency shifting while ensuring continuous Fourier-space coverage. The spatial distribution of these patterns is modulated by the illumination angle, as predicted by numerical simulation (Figure S2). The near-field distribution surrounding the nanoholes was experimentally confirmed by cathodoluminescence characterisation (Supplementary Note 1, Figure S3). The angle-dependent fields provide the structured illumination required to shift high spatial frequency information into the detectable bandwidth.

**Figure 1.**
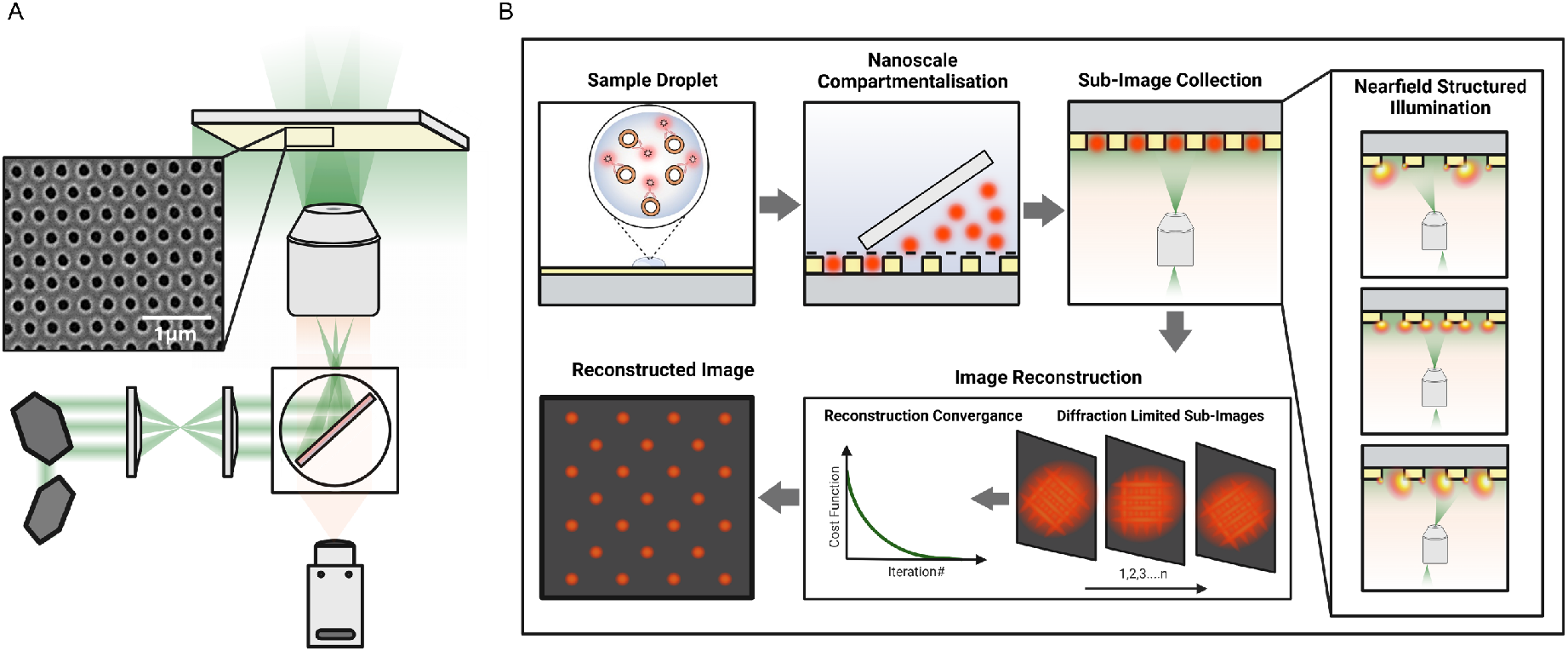
Schematic of the NHA-assisted super-resolved single sEV assay. **A** Optical setup for the near-field assisted SIM platform. The system integrates a plasmonic NHA into the SIM light path to generate on-chip sub-diffraction illumination patterns. A galvanometric mirror system directs the laser beam across the back focal plane of a high-numerical-aperture objective lens, enabling wide-field illumination with angular control. This configuration provides continuous adjustment of the azimuthal angle (ϕ) from 0° to 360° and supports polar incident angles (θ) up to ±56°. When the NHA is illuminated near its dipole resonance, these angular sweeps produce structured near-field illumination patterns, which are essential for SIM. Inset: Scanning electron microscope image of the fabricated NHA. **B** Workflow of the super-resolved single sEV assay. A 20 μL sample droplet containing pre-labelled sEVs is deposited on the NHA surface, where vesicles are discretely confined within individual nanoholes via capillary-assisted compartmentalisation^22^. Near-field structured illumination is generated by exciting plasmonic modes at varying incident angles. Each angle generates a distinct near-field distribution, resulting in a corresponding sub-image of the fluorescently labelled sEVs. Tuning the incident angle effectively shifts the illumination phase, while varying the azimuthal angle alters the pattern orientation, analogous to phase stepping and pattern rotation in conventional SIM. These angle-dependent sub-images are computationally reconstructed using the Blind-SIM algorithm to yield a super-resolved image.

Unlike previous plasmonic-assisted SIM platforms that primarily function as structured illumination substrates^17,18,19^, the NHA architecture introduced here serves a dual-function interface. Each nanohole acts as a nanoscale compartment for particle localisation, while the periodic plasmonic lattice simultaneously generates the near-field illumination patterns required for super-resolution reconstruction. To achieve isotropic reconstruction, raw sub-image stacks comprising 9, 12, or 24 raw images were acquired by systematically tuning the incident angle (*θ*) and azimuthal angle (*φ*). These variations generate distinct near-field patterns across the NHA, encoding complementary high-spatial-frequency information from the same confined particles, which is subsequently reconstructed using a blind-SIM algorithm^20,21^. Notably, this periodic nanostructure design enables highly efficient spatial-frequency encoding, requiring significantly fewer sub-images than stochastic, speckle-based metasurface SIM approaches^17,21^. Numerical simulations comparing NHAs to nanopillar arrays (NPAs) confirmed that although the NHAs support a hybrid of localised and propagating plasmon modes, this modal complexity does not compromise the fidelity of resolution recovery (Figure S4). Finally, control experiments of imaging 100 nm fluorescent polystyrene nanoparticles on both glass and planar gold failed to yield any super-resolution, confirming that the engineered NHA architecture is required for resolution enhancement (Supplementary note 2, Figure S5).

Figure 1B illustrates the single sEV assay workflow. sEVs pre-labelled with fluorescent antibodies are deposited on the NHA surface via capillary-assisted scraping^22^. This meniscus-guided assembly stochastically anchors vesicles into nanoholes for single-particle confinement. The EV-loaded NHA is immersed in 1× Phosphate-Buffered Saline (PBS) and imaged using a customised SIM setup (Figure 1A and Figure S6). To generate the structured illumination pattern, a linearly polarised laser (λ = 561 nm) is steered by a pair of galvanometric mirrors through a 4*f* optical system and directed onto the NHA via a high-numerical-aperture objective lens. Azimuthal (0°–360°) and polar (±56°) angular tuning at the back focal plane selectively excites near-field plasmonic modes, shifting the phase and orientation of the sub-diffraction patterns. Sub-images are acquired and reconstructed using a blind-SIM algorithm to obtain super-resolved images for quantifying sEV counts^20,21^.

### Validation of nanoparticle confinement and spatial resolution via fluorescent nanoparticle standards

To demonstrate the dual functionality of NHAs for both single-particle confinement and super-resolution interrogation, we first validated the platform using 100 nm fluorescent polystyrene nanoparticle standards. A 20 μL sample (1×10^12^ particles/mL in water) was applied to the NHA surface via capillary-assisted scraping. Particle confinement is driven by evaporation-induced capillary assembly at the receding contact line, where convective flow transports particles toward the liquid interface. As the meniscus traverses the NHA, local pinning over the nanostructured topography increases particle residence time and promotes trapping within individual nanoholes, consistent with established convective assembly processes on patterned substrates^12,13,22^. Because this process is governed primarily by particle flux to the receding interface rather than bulk concentration, evaporation-induced interfacial enrichment can sustain confinement even at low particle concentrations^23^. Scanning electron microscopy (SEM) confirms precise nanoscale compartmentalisation of particles within individual nanoholes (Figure 2Ai, 2Aii and S7). Figure S8 provides a correlative validation of nanoparticle confinement from the same particles under SEM, diffraction-limited and NHA-SIM images. Importantly, particle localisation at the NHA does not require complete physical insertion into the 90 nm apertures. Control experiments using nominally 200 nm polystyrene nanoparticles demonstrated that these larger targets preferentially anchor at the nanohole rims (Figure S9), validating an aperture-mediated confinement that operates independently of absolute particle size, rendering the platform resilient to the natural size heterogeneity of sEVs.

**Figure 2.**
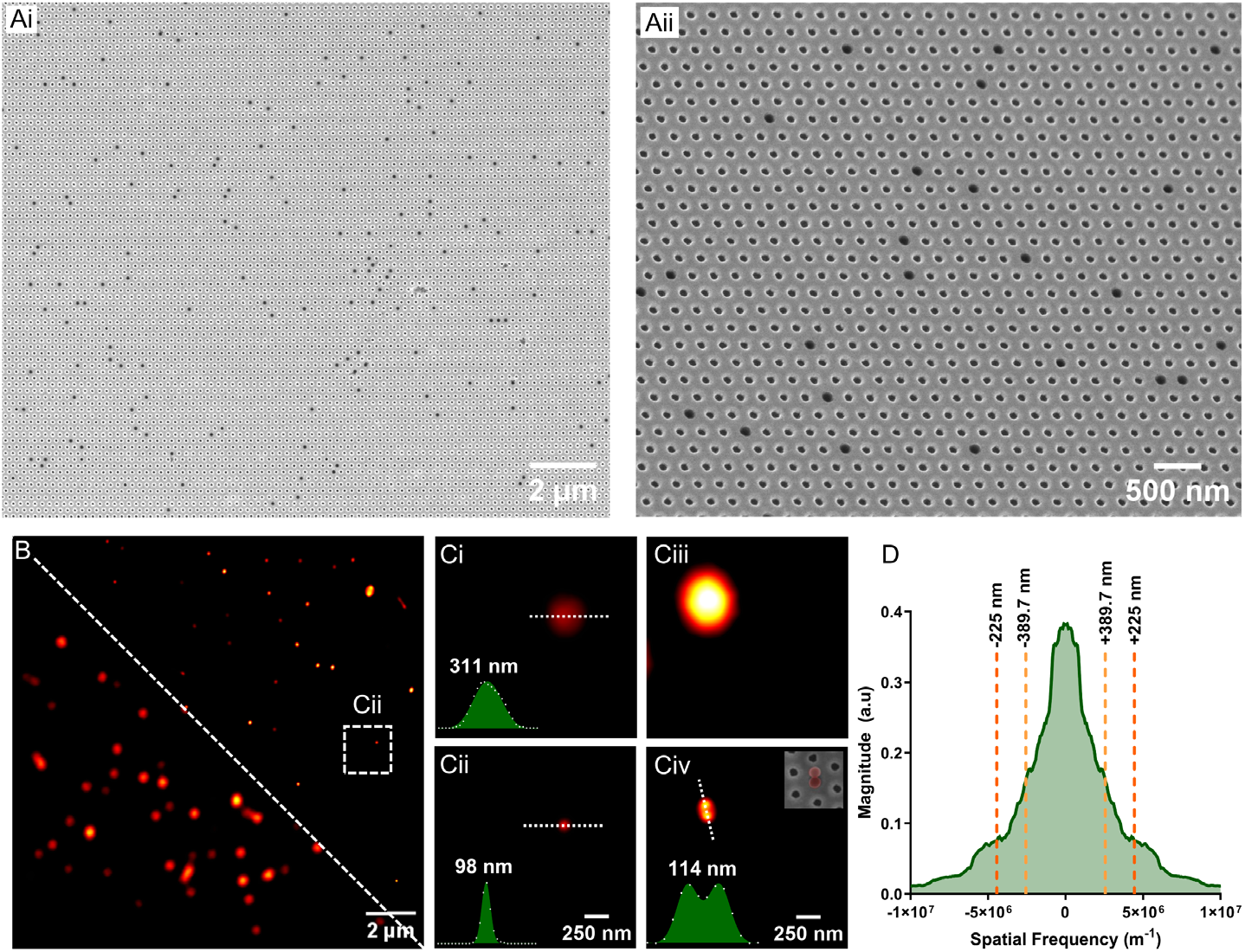
Validation of nanoparticle confinement and spatial resolution via fluorescent standards. **Ai** Low magnification SEM image showing compartmentalised 100 nm polystyrene nanoparticles within individual nanoholes. **Aii** Higher-magnification, 40°-titled SEM image highlighting the nanoscale confinement of individual particles. **B** Diffraction-limited widefield fluorescence image (left) and the corresponding NHA-SIM super-resolved image (right). White dashed boxes denote ROIs for magnified views in ci and cii. **Ci** Diffraction-limited image of a single particle showing a full width at half maximum (FWHM) of 311 nm. **Cii** NHA-SIM image of the same particle with a FWHM of 98 nm. **Ciii** Diffraction-limited image failing to resolve two nanoparticles compartmentalised within a single nanohole. **Civ** Corresponding NHA-SIM image of the same dimer, successfully resolved with a centre-to-centre separation of 114 nm. Inset: SEM image of the same dimer. **D** Fourier domain analysis of the reconstructed image over the full field of view (16 µm). The 1D line profile reveals well-defined peaks matching the spatial frequencies of the nanohole pitch (225 nm) and its 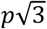 harmonic (389.7 nm).

**Figure 3.**
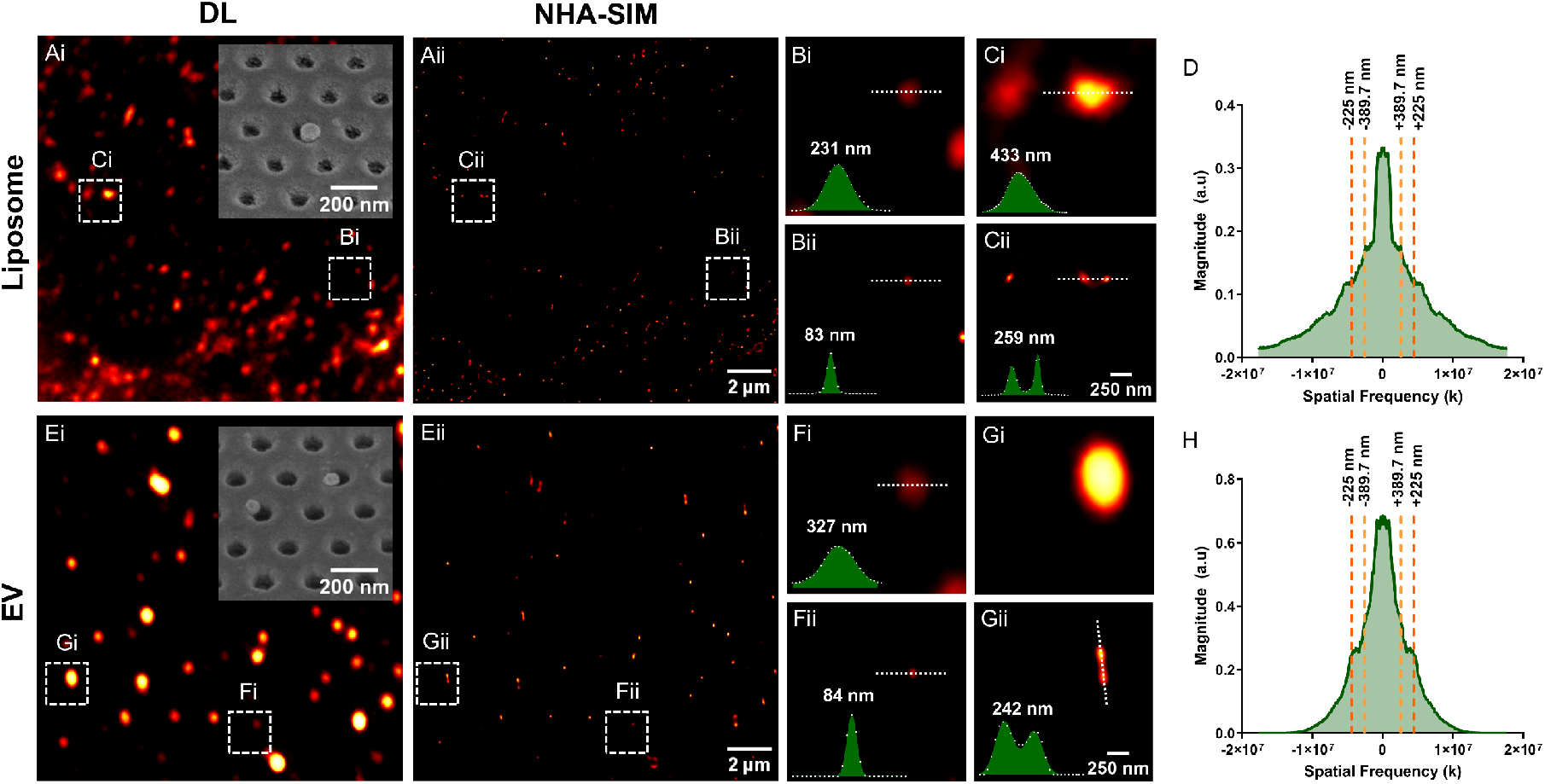
Validation of particle confinement and spatial resolution via fluorescent liposome and sEVs. **Ai** Diffraction-limited (DL) image and **Aii** the corresponding NHA-SIM image of compartmentalised 100 nm fluorescent liposomes. White dashed boxes denote ROIs for magnified views. Insets: SEM images of liposomes within nanoholes. **Bi** Diffraction-limited and **Bii** NHA-SIM images of a single liposome showing FWHMs of 231 nm and 83 nm, respectively. **Ci** Diffraction-limited and **Cii** NHA-SIM image of liposome dimers, resolving a centre-to-centre separation of 259 nm. **D** 1D Fourier transform of the reconstructed image of liposomes showing peaks matching the spatial frequencies of the nanohole lattice. **Ei** Diffraction-limited and **Eii** NHA-SIM images of compartmentalised 100 nm m-Cherry sEVs. White dashed boxes denote ROIs for magnified views. Insets: SEM images of sEVs within the nanohole. **Fi** Diffraction-limited and **Fii** NHA-SIM images of a single sEVs showing FWHMs of 327 nm and 84 nm, respectively. **Gi** Diffraction-limited and **Gii** NHA-SIM images of sEV dimers, resolving a centre-to-centre separation of 242 nm. **H** 1D Fourier transform of the reconstructed image of sEVs with peaks matching the spatial frequencies of the nanohole lattice.

Following physical validation, we next assess the optical performance of the proposed imaging modality using these compartmentalised particles. Figure 2B compares a diffraction-limited image (left) with a reconstructed NHA-SIM image (right). For a single confined nanoparticle, the plasmonic nanohole array structured illumination microscopy (NHA-SIM) image yields a full-width at half-maximum (FWHM) of 98 nm, an approximate 3-fold reduction compared to the 311 nm diffraction-limited FWHM (Figure 2Ci, Cii). Although the platform predominantly achieves single-particle occupancy, occasional instances of two nanoparticles per hole occur (Figure 2Ciii, iv). NHA-SIM successfully resolve these dimers, revealing a centre-to-centre separation of 114 nm (Figure 2Civ).

To further confirm particle confinement, we performed a Fourier transform of the super-resolved fluorescence image (Figure 2Bi). The one-dimensional line profile extracted from the Fourier spectrum along the principal lattice direction (Figure 2D) exhibits peaks at spatial frequencies corresponding to the NHAs pitch (225 nm) and its 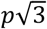 harmonic (389.7 nm). The emergence of these lattice-matched frequency components indicates that the fluorescence emission distribution inherits the underlying NHA architecture. The relatively low amplitude of these peaks is attributed to the dominant background signal (unfilled nanoholes), which reduces the overall contrast in the Fourier domain. Critically, the fixed positioning of each particle ensures continuous interaction with the structured near field, enabling consistent spatial-frequency mixing that shifts sub-diffraction features into the optical system’s resolvable bandwidth.

### Validation of nanoparticle confinement and spatial resolution via fluorescent liposomes and sEVs

Following the validation with polystyrene standards, we evaluated the platform’s performance using biologically relevant lipid nanoparticles, specifically artificial liposomes and cell-derived sEVs. Unlike solid fluorescent polystyrene nanoparticles, lipid nanoparticles present a more complex imaging target due to heterogeneous fluorophore density, varied membrane compositions, and delicate morphologies.

Fluorescent liposomes (labelled with DiL membrane dye) and sEVs (derived from genetically engineered human embryonic kidney cells HEK293 cells expressing mCherry-CD9) were sourced from commercial suppliers (Table S1). DiL and mCherry were selected to align with the system’s excitation and emission specifications. Nanoparticle tracking analysis (NTA) confirmed a median hydrodynamic diameter of 108 nm for liposomes and 120.4 nm for sEVs (Supplementary Note 3), with vesicle morphology further verified by cryo-electron microscopy (Figure 4C). Liposomes (1 × 10^12^ particles/mL) and sEVs (1 × 10^9^ particles/mL) were deposited onto the NHA using the previously described capillary-assisted scraping method.

**Figure 4.**
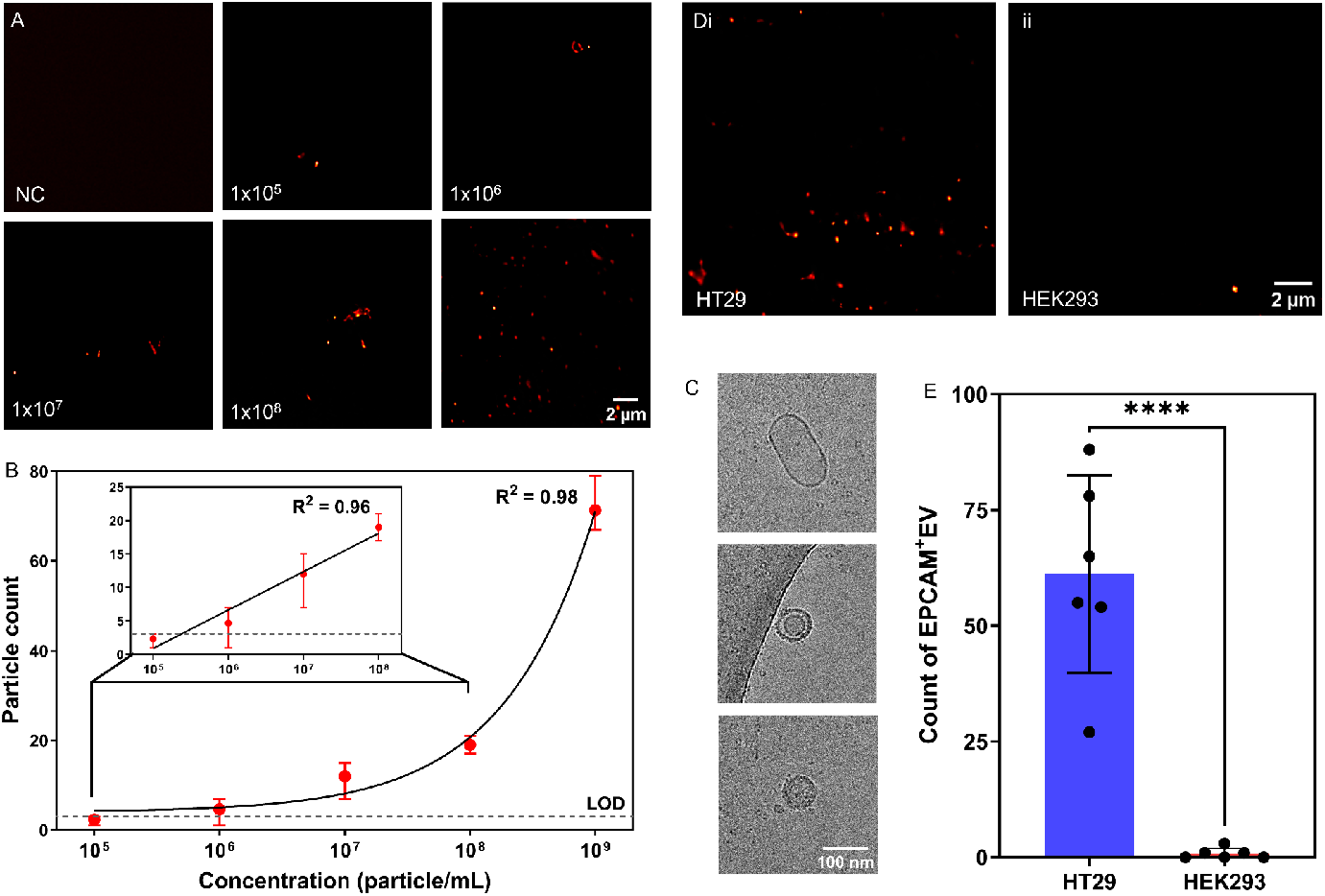
Analytical performance validation of the super-resolved single sEV assay. **A** Representative NHA-SIM images of mCherry-labelled HEK293 sEVs across a serial dilution series (1 × 10^5^ to 1 × 10^9^ particles/mL), alongside a negative control (NC) of 1 × 10^9^ particles/mL unlabelled HEK293-derived sEVs. **B** Calibration curve showing particle count (mean ± range) as a function of EV concentration (n = 3), with a 4PL fitting. The LOD is indicated by the dashed line, corresponding to 143 sEVs/μL. The inset shows the linear response regime (R^2^ = 0.96). **C** Cryo-electron microscopy images of mCherry-labelled sEVs confirming vesicle morphology. **D** Representative NHA-SIM images of EpCAM^+^ sEVs from **Di** HT29 cells and **Dii** HEK293 cells. **E** Quantitative analysis of EpCAM^+^ EV counts from HT29 and HEK293 cells (n = 6, ****p < 0.0001).

Comparative diffraction-limited and NHA-SIM images are presented in Figure 3Ai–Cii (liposomes) and 3Ei–Gii (sEVs). For liposomes, the FWHM decreased from 231 nm to 83 nm, representing an approximate 2.8-fold improvement (Figure 3Bi–Bii). For sEVs, the FWHM was reduced from 327 nm to 84 nm, an approximate 3.9-fold improvement over the diffraction limit (Figure 3Fi–Fii). The measured FWHM differs from the physical particle size; this discrepancy arises from the combined effects of iterative Blind-SIM reconstruction and highly confined near-field illumination, which preferentially enhance localised fluorescence regions within the spherical particles, an effect consistent with related metasurface-based SIM studies^18,24^. This effect contributes to the small measured FWHM values observed in the reconstructed images.

The platform’s dual functionality enables the resolution of closely spaced particles within adjacent nanoholes. Reconstruction of liposome and EV dimers (Figure 3Ci–Cii and 3Gi–Gii) reveals distinct intensity maxima separated by 259 nm (liposomes) and 242 nm (sEVs). These values closely match the 225 nm pitch, confirming geometry-defined confinement within neighbouring nanoholes. The slight deviations observed are likely due to off-centre particle positioning within the nanoholes, potentially arising from the slightly tapered nanohole sidewalls produced during fabrication. In addition, non-uniform excitation arising from localised near-field enhancement and evanescent-field illumination may influence the reconstructed particle intensity distribution and apparent localisation, contributing to small deviations in the measured centre-to-centre distances compared to the NHAs periodicity. Furthermore, in biological particles such as EVs, variations in fluorophore density, fluorophore distribution, and particle morphology may further contribute to this variability.

SEM imaging further supports particle localisation within the nanoholes (Figure 3Ai and 3Ei insets, Figure S11). However, the inherent fragility of biological structures can lead to particle loss during the sample preparation (fixation and drying) and imaging process (electron-beam exposure) required for SEM (Supplementary Note 4), resulting in lower observed densities compared to fluorescence imaging. Furthermore, Fourier-domain analysis of both liposomes and sEVs (Figure 3D and 3H) showed peaks matching the nanohole lattice pitch, further reinforcing the evidence of architectural confinement. Finally, benchmarking against a commercial TIRF system using open-source decorrelation analysis^25^ showed a 3.5-fold resolution improvement, confirming that the resolution enhancement is intrinsic to the NHA-SIM interface rather than system-specific artefacts (Supplementary Note 5). The larger resolution improvement measured by decorrelation analysis likely arises because it captures the combined effects of near-field plasmonic confinement and iterative Blind-SIM reconstruction, rather than only the idealised spatial frequency gain predicted from the illumination pattern periodicity.

### Single EV assay and analytical performance validation

The analytical sensitivity and specificity of the NHA-SIM platform were evaluated through direct quantification of single sEVs from the super-resolved images. Figure 4A displays representative NHA-SIM images of m-Cherry sEVs across a five-log concentration range (1 × 10^5^ to 1 × 10^9^ EVs mL^−1^), alongside a negative control of 1 × 10^9^ particles/mL unlabelled HEK293-derived sEVs. The resulting calibration curve (Figure 4B) demonstrates a concentration-dependent increase in particle counts, as characterised by a four-parameter logistic (4PL) fit (R^2^ = 0.98), a feature characteristic of quantitative bioassays. In the low-concentration regime (10^5^–10^8^ particles/mL), the response is highly linear (R^2^ = 0.96), ensuring reliable quantification of single EVs in this dynamic range. No measurable fluorescent particle signal was observed in the negative control (NC), with images exhibiting only background-level intensity and no detectable localised fluorescent emission. Consequently, analytical noise was estimated from the regression residuals according to established analytical validation guidelines^26^. The LOD was determined via a weighted least-squares fit to this linear region (LOD = 3σ_γ/x_ / m), where σ_γ/x_ is the residual standard deviation of the regression and m is the slope. This yielded a signal threshold of approximately 3 counts per field of view (FOV). This corresponds to an LOD of 1.43 × 10^5^ particles/mL (143 sEVs/μL), which is highly comparable to that of state-of-the-art single-EV counting technologies (Table S4). Although diffraction-limited imaging captured the overall concentration-dependent trend, it exhibited significantly compromised counting fidelity, characterised by poor linearity (R^2^ = 0.60) and marked deviations from a monotonic response (Figure S13). Crucially, because the diffraction-limited readout maintains a predictable tracking capability at lower concentrations where particle density is sparse, the dramatic drop in linearity points directly to a density-dependent resolution-coincidence bottleneck. As spatial crowding increases, standard optics inevitably merge closely spaced features into a single signal. NHA-SIM cleanly overcomes this packing-density limitation, leveraging near-field super-resolution to accurately resolve individual sub-wavelength targets and restore true digital linearity across the entire concentration range.

To assess analytical specificity, we compared sEVs derived from HT29 (human colorectal adenocarcinoma) and HEK293 cell lines. HT29-derived sEVs are known to express significantly higher levels of the surface marker EpCAM than HEK293-derived vesicles, as validated in our previous studies using both bulk and single-EV analyses^27,28^. Before depositing to the NHA, sEVs were incubated with Alexa Fluor 549 conjugated anti-EpCAM antibody, with excess free fluorescent antibody removed by size-exclusion chromatography following our previous protocol^28^. Figure 4Di–ii presents representative NHA-SIM images, showing a markedly higher abundance of EpCAM^+^ sEVs in the HT29 samples. Quantitative analysis further confirmed this difference, with an average of 61.2 ± 21.3 particles per FOV for HT29 versus 0.8 ± 1.2 particles per FOV for HEK293 (Figure 4E). These results validate the platform’s ability to distinguish sEV subpopulations based on specific protein expression.

## Discussion

The NHA-SIM architecture bridges the gap between digital bioassays and super-resolution imaging by integrating plasmonic NHAs with near-field SIM. Unlike conventional digital assays that rely on stochastic Poisson loading, this platform localises analytes within a geometry-defined nanoscale array while simultaneously interrogating them with sub-diffraction near-field illumination patterns. By physically registering particles to discrete nanohole positions, the NHA provides a spatially defined readout framework that reduces coincidence events; within this architecture, the near-field assisted SIM actively resolves individual sub-wavelength particles, preventing multi-occupancy signal merging and improving digital counting fidelity. This dual functionality enables both discrete single-particle confinement and super-resolved optical readout, eliminating the need for droplet encapsulation, enzymatic amplification, or complex optical architectures. Furthermore, the platform is fully compatible with standard fluorescence-based detection, scalable nanofabrication and passive sample preparation. The broadband plasmonic response of gold nanoholes additionally supports multicolour fluorescence imaging, offering a path towards multiplexed biomarker detection and marker colocalization analysis from single EVs.

Particle occupancy within the NHA remained intrinsically sparse across all tested concentrations, suggesting that loading is highly modulated by local capture dynamics, including surface charge and capillary forces, rather than concentration alone. While this sparsity ensures single-particle isolation and minimises signal overlap, it reduces the fraction of active sensing sites. Future strategy could improve occupancy by enhancing particle–nanohole interactions through surface functionalisation, optimising nanohole geometry and periodicity, or introducing an incubation period prior to scraping to allow for particle settling^29,30^ . In addition, expanding the FOV through larger arrays would increase the total number of measurable events, enhancing statistical sampling and detection sensitivity. Integrating dielectrophoresis within a microfluidic environment could further guide particles toward uniform trapping regions, stabilising the LOD and improving reproducibility^31^.

Future design directions may include alternative plasmonic materials, such as silver-based architectures for enhanced near-field confinement, to further improve the optical resolution^32^, or chirped NHAs with spatially graded geometries for size-selective sorting of vesicles. Although this study utilised purified vesicle samples, translation to clinical specimens represents a natural next step. The challenges posed by complex biofluids can be mitigated by combining NHA-SIM with established EV isolation workflows. Uniquely, the defined physical architecture of the NHA provides opportunities for further image enhancement through post-processing approaches such as spatial filtering, deconvolution, and mask-guided analysis, which can help suppress background contributions, improve image quality and particle enumeration. However, in this work, the analysis was intentionally focused on the blind-SIM reconstruction workflow to evaluate the intrinsic performance of the proposed imaging platform. Ultimately, by unifying geometry-defined particle registration with position-deterministic near-field super-resolution imaging, the NHA-SIM platform successfully overcomes the fundamental resolution-coincidence trade-off of conventional digital assays, establishing a robust framework for high-fidelity single-vesicle analysis in complex biological environments.

## Methods

### Fabrication of Plasmonic Nanohole Arrays

NHAs were fabricated in accordance with our previously established protocol at the Australian National Fabrication Facility (ANFF) – NSW Node (Figure S14). A 1mm thick fused silica wafer (Micro Materials Pty Ltd) was used as the substrate. A 10 nm titanium adhesion layer followed by an 80 nm gold film was deposited via thermal physical vapour deposition (Lesker PVD75 e-beam evaporator) at a deposition rate of 0.5 Å s^−1^. Nanohole patterns were defined using electron beam lithography (EBL; Raith 150-Two) with AR-P 6200.09 positive-tone resist (Allresist). Pattern transfer into the gold film was achieved via argon ion beam etching (NANOQUEST I Ion Beam Etcher).

### SAMPLE deposition with capillary-assisted scraping

A scraping-assisted, meniscus-guided assembly method was used to load polystyrene beads, liposomes, and sEVs into the NHAs. The particle concentrations were determined by NTA before deposition (Table S3). A 20 µL aliquot of sample was pipetted onto the substrate, and the droplet was slowly translated across the surface using a cut piece of PMMA. The meniscus, pinned at the PMMA edge, was swept across the substrate, directing particles at the meniscus–substrate interface into the nanoholes. The PMMA chip was held at a shallow angle (∼15–20°) to maintain a stable contact line and promote consistent particle transport into the array.

### NHA-SIM imaging and image reconstruction

A 561 nm laser was passed through a Fourier filtering beam-cleaner to enhance beam quality before being collimated to a beam size of 1–2 mm^2^. The incident angle of illumination was controlled via a two-dimensional (2D) galvanometric scanning mirror system (Thorlabs) in conjunction with a high-numerical-aperture (NA) 4f optical system, allowing angular tuning within a ±56° range. To minimise vibrations and enhance optical stability, the entire setup was mounted on a floating optical table.

For NHA-SIM reconstruction, raw sub-image stacks were acquired under defined incident and azimuthal illumination angles. The polystyrene nanoparticle images shown in Figure 2 were reconstructed from 12 sub-images acquired at a fixed incident angle of θ = 40°, with the azimuthal angle φ varied from 0° to 330° in 30° increments. For the liposome reconstruction in Figure 3, 24 sub-images were acquired using two incident angles, θ = 30° and θ = 40°, with 12 azimuthal angles collected for each incident angle from φ = 0° to 330° in 30° increments. For the extracellular vesicle reconstructions in Figure 3, 9 sub-images were acquired using three incident angles, θ = ™40°, 0°, and 40°, combined with three azimuthal angles, φ = 0°, 60°, and 120°. The limit-of-detection and EpCAM-specific EV reconstructions shown in Figure 4 were acquired using the same 9-image angular acquisition scheme. Fluorescent emission from the sample was collected using a Nikon 100× oil immersion objective (NA 1.25). After filtering out residual excitation wavelengths, the signal was directed through a tube lens and captured by a CMOS camera (Dhyana 400BSI V3).

A custom LabVIEW program was implemented to manage both experimental synchronisation and voltage control. The galvanometric mirrors were modulated through an analogue voltage signal generated within this program, ensuring precise angular adjustments. While the camera was interfaced through proprietary software, its digital trigger signal for frame capture was also controlled within the same LabVIEW environment, ensuring full synchronisation of SIM image sequences.

All image super-resolution reconstructions were conducted in MATLAB 2023a. Super-resolved images were obtained using a blind Structured Illumination Microscopy (blind-SIM) algorithm^20,21^, which does not require prior knowledge of the illumination patterns. Instead, both the object and the near-field illumination distributions are treated as unknowns and jointly recovered in real space through an iterative cost-minimisation strategy.

### EV labelling with fluorescent antibody

For EV labelling, we used the protocol developed in our previous work. Specifically, 30 μL of 0.25mg/mL AF549-conjugated anti-EpCAM antibody was incubated with 300 μL HEK293 EVs (1.3 × 10^10^ particles/mL) and HT29 EVs (4.1 × 10^10^ particles/mL) at room temperature in the dark for 2 hours. After labelling, all samples were topped up to a final volume of 1 mL with filtered 1×PBS and loaded onto the SEC column for unbound dye removal. Post-labelling samples were subsequently characterised using NTA. (Supplementary Note 3) These samples were then directly used for deposition onto NHA substrates.

### Single EV assay quantification and data analysis

All data were analysed and visualised using GraphPad Prism (version 10.4.2) and MATLAB (MathWorks, R2023b). MATLAB was used for image-based analysis, including line profile extraction, Gaussian fitting, and Fourier transform analysis. GraphPad Prism was used for data visualisation and statistical analysis. Particle enumeration was performed using a peak-based detection workflow in Fiji/ImageJ, adapted from local-maxima-based single-particle counting approaches. Detected maxima were exported as calibrated x–y coordinates and manually verified against the original unprocessed reconstructions to remove false positives and avoid duplicate counting. This workflow is consistent with previously published single-particle imaging approaches that use local-maxima detection for particle enumeration^33^.

For calibration curve analysis, particle counts were extracted from NHA-SIM images for each EV concentration, and the mean particle count was calculated from 3 FOVs per concentration. Data are presented as mean ± standard deviation, with error bars representing the standard deviation between FOVs. Calibration curves were fitted using a four-parameter logistic (4PL) regression model. The linear dynamic range was determined by linear regression analysis of the linear portion of the calibration curve.

In the absence of a measurable signal from the blank control, the analytical noise was estimated from the residual standard deviation of the regression (σ_γ_/_x_) in accordance with established analytical validation guidelines^26^. The LOD was subsequently calculated from the weighted least-squares linear fit using the relationship LOD = 3σ_γ_/_x_ / m, where m represents the slope of the fitted regression line. Statistical comparisons between EpCAM expression levels of HT29-derived EVs and HEK293-derived EVs were performed using an unpaired two-tailed t-test based on particle counts obtained from 6 FOVs per group.

## Acknowledgements

This work was supported by the Australian Research Council Discovery Early Career Researcher Award (DE240100321) awarded to Y. Z., and the Australian Research Council Discovery Project (DP260101407) awarded to P. J. R. We also acknowledge Alexander G. D. MacIntosh and Adam Stewart from the School of Physics, UNSW Sydney, for their valuable discussion regarding imaging and data analysis. We acknowledge the Australian National Fabrication Facility (ANFF) NSW Node for training and support in fabrication processes. We acknowledge Microscopy Australia and the Victor Chang Cardiac Research Institute Innovation Centre for use of SEM, TEM and Cryo-TEM facilities at the Electron Microscope Unit (EMU) within the Mark Wainwright Analytical Centre (MWAC) at UNSW Sydney.

## Author Information

Authors and Affiliations

**School of Biomedical Engineering, University of Technology Sydney, NSW, Australia**

Anthony J. El-Helou, Yiting Liu, Fatemeh Khosravi, Chaohao Chen, Ying Zhu

**School of Physics, UNSW Sydney, NSW, Australia**

Peter J. Reece

**Electron Microscope Unit, UNSW Sydney, NSW, Australia**

Carol H.W. Yan, Mark Lockrey, Juanfang Ruan

**Department of Electrical and Computer Engineering, University of California, San Diego, CA, USA**

Zhaowei Liu

**School of Clinical Medicine, UNSW Sydney, NSW, Australia**

Ying Zhu

## Author contributions

Y. Z. and P. J. R. conceived, designed, and supervised the research. A.J.E. fabricated the plasmonic metasurfaces, built the optical setup, carried out the experiments, performed the data analysis, and drafted the manuscript. Y. L. prepared the biological samples and performed NTA measurements. F.K. assisted with sample preparation and figure preparation. C. C. provided feedback on experimental design and data interpretation. C. H.W. Y. performed sample preparation and fixation prior to SEM imaging. M.L. performed the cathodoluminescence measurements. J. R. conducted the cryo-EM imaging. Z. L. provided expert guidance and critical feedback. All authors contributed to the writing and revision of the manuscript.

Correspondence and requests for materials should be addressed to Ying Zhu.

## Data availability

Source data are included in the Supplementary Information. Additional information supporting this study can be obtained from the corresponding author upon reasonable request.

