## Supplementary Information for "Super-Resolved Single Small Extracellular Vesicle Assay enabled by a Plasmonic Nanohole Array"

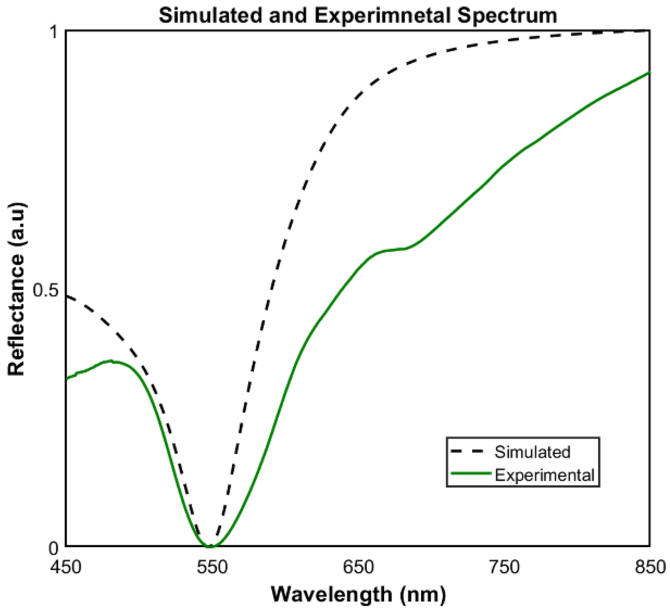


**Figure S1: Comparison of simulated and experimental spectrum**. Simulated (dashed line) and experimental (solid line) reflectance spectra of NHA engineered with a 90 nm hole diameter and 225 nm periodicity, showing close spectral agreement.

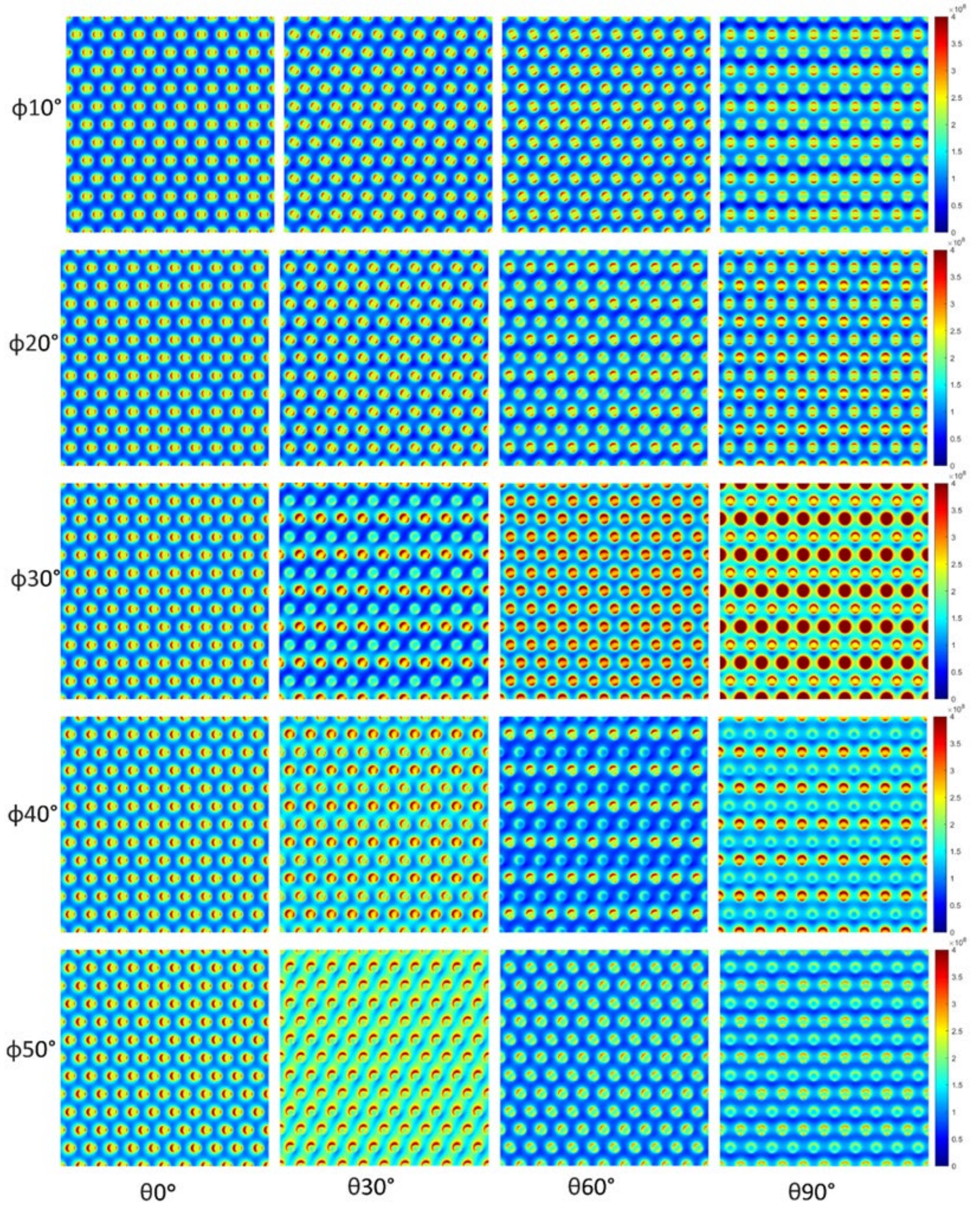


**Figure S2: Simulated structured near-field intensity distributions of the hexagonal nanohole array at resonance for different incident angles and azimuthal orientations**. Rows correspond to increasing incident angle, ϕ = 10°–50°, while columns correspond to azimuthal orientation, θ = 0°, 30°, 60° and 90°. The colour scale represents the simulated near-field intensity, highlighting the angle- and orientation-dependent modulation of the resonant field patterns. These variations demonstrate the ability of the NHA to generate diverse structured near-field distributions with spatial frequency content relevant for SIM-based super-resolution imaging.

**Supplementary Note 1: Cathodoluminescence Characterisation of Patterned Gold Films**

To evaluate film quality and confirm plasmonic near-field generation necessary for structured illumination, cathodoluminescence (CL) imaging was performed using a Nova SEM 450 equipped with a CL module. Direct CL characterisation of the NHA geometry used for NHA-SIM was challenging because CL measurements are conducted in a vacuum. In contrast, the imaging system operates in solution, resulting in a blue shift of the plasmonic resonance due to the lower refractive index of the surrounding medium. This shift moves the resonance closer to the gold interband absorption region (~532–561 nm), where CL signal detection becomes difficult due to both reduced plasmonic emission and the limited sensitivity range of the CL CCD detector. In addition, $SiO_2$ substrates can exhibit visible emission associated with self-trapped excitons under electron irradiation[1], which spectrally overlaps with the NHA-SIM resonance region and complicates near-field interpretation. Consequently, larger-pitch NHAs (P580) fabricated on suspended silicon nitride (SiN) membranes were employed to shift the plasmonic response toward longer wavelengths, enabling clearer visualisation of the localised near-field behaviour and qualitative assessment of the quality of the fabricated gold film.

Figure S3A shows an SEM image of the fabricated NHA, with the dashed red region indicating the CL acquisition area. Spatially resolved CL intensity maps are presented in Figure S3Bi–iii for the spectral ranges 500–550 nm, 700–750 nm, and 800–950 nm, respectively. A pronounced ring-like emission pattern surrounding the nanoholes is observed specifically within the 700–750 nm range, consistent with localised plasmonic excitation at the nanohole edges. The ring-like symmetry arises from the unpolarised nature of the CL measurement, which averages emission over all dipole orientations. Following background subtraction (Figure S3Ci–iii), the ring-like features become more clearly defined, confirming localised near-field enhancement at the nanohole boundaries. To further validate this behaviour, CL spectra extracted from the nanohole edge, aperture centre, and inter-hole region are shown in Figure S3D. Spectra obtained at the nanohole perimeter exhibit a clear resonance peak within the 700–750 nm range. In contrast, spectra between nanoholes show no corresponding enhancement, confirming that the observed emission originates from localised plasmonic modes concentrated at the nanohole edges.

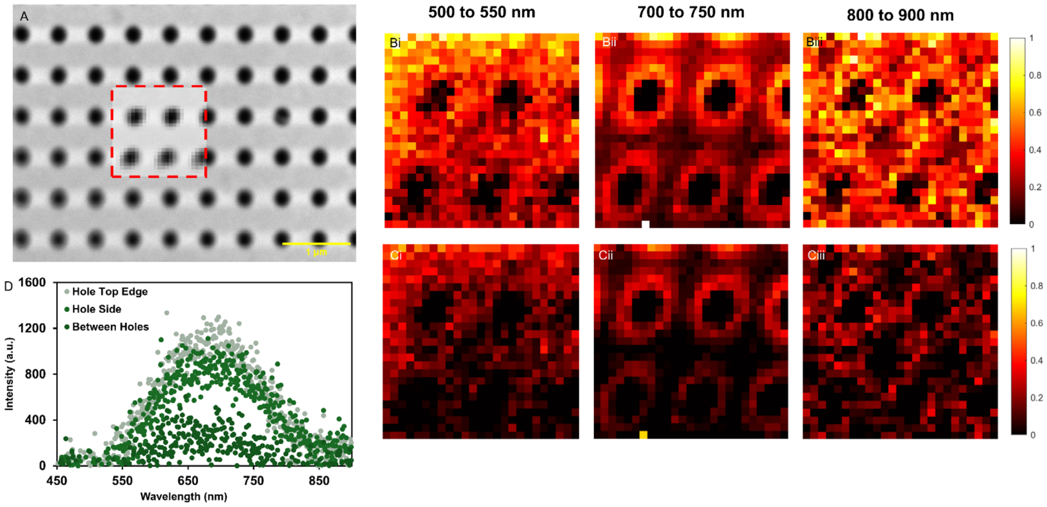

**Figure S3: Cathodoluminescence (CL) characterisation of the gold nanohole array. A** SEM image of the nanohole array, with the dashed red region indicating the area analysed by CL mapping. **Bi–iii**, CL intensity maps acquired over three spectral windows: 500–550 nm, 700–750 nm, and 800–950 nm. A distinct ring-like emission pattern is observed around the nanoholes within the 700–750 nm band, indicating wavelength-selective plasmonic excitation. **Ci–iii.** Corresponding background-subtracted CL maps, which enhance the visibility of the localised emission features at the nanohole edges. **D.** CL spectra extracted from different spatial locations, including the nanohole edge, within the aperture, and between nanoholes. A pronounced spectral peak is observed at the nanohole perimeter within the resonance band (700–750 nm), confirming that the emission originates from localised plasmonic modes.

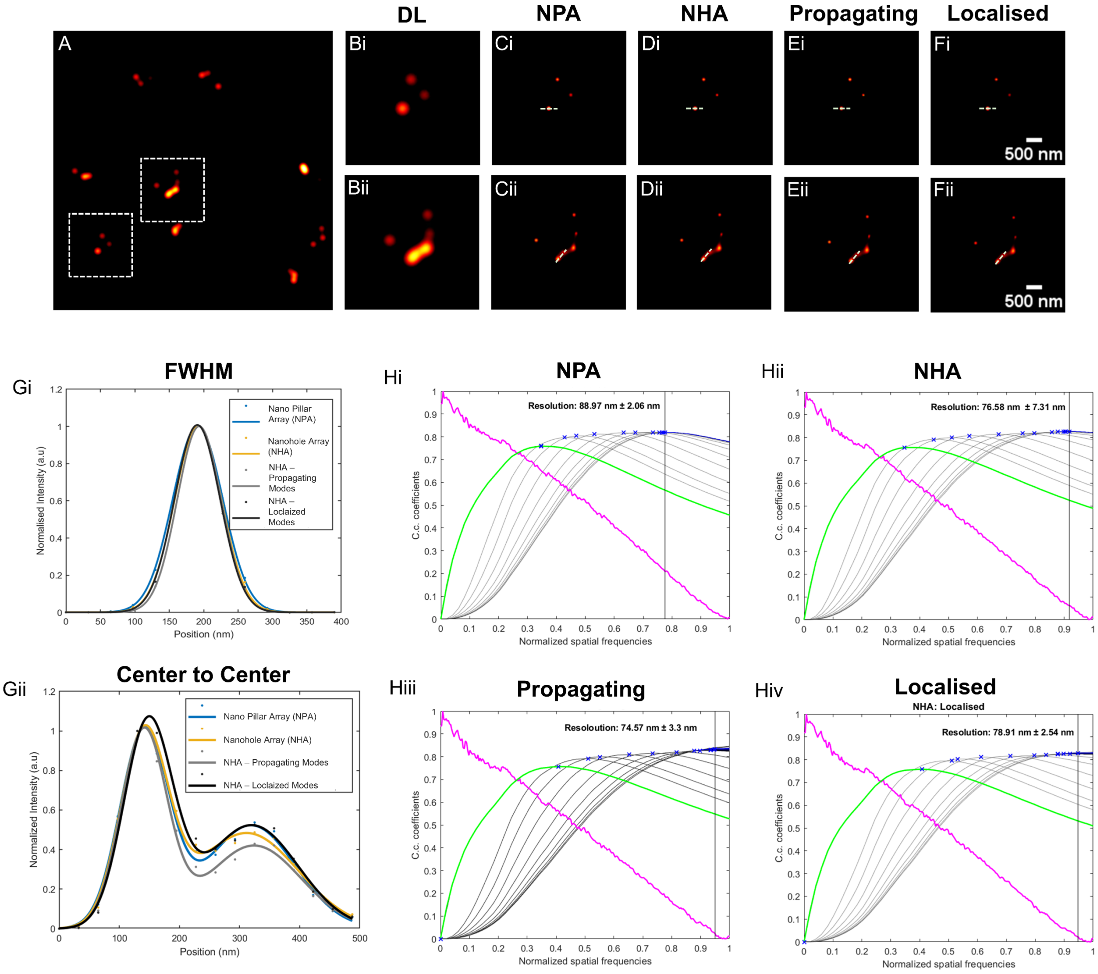


**Figure S4: Comparative analysis of reconstruction resolution for plasmonic nanopillar array (NPA) and NHA. A** Simulated Image. **Bi–ii** Diffraction-limited (DL) images of selected regions of interest, denoted by the white dashed boxes. **Ci–ii** NPA SIM reconstructions. **Di–ii** NHA-SIM reconstructions. **Ei–ii** Reconstructions using propagating mode subset. **Fi–ii** Reconstructions using localised mode subset. **Gi–ii** Line intensity profiles corresponding to the indicated regions of interest, showing comparable peak narrowing and feature separation across all configurations. **Hi–iv** Decorrelation analysis[2] used to estimate resolution, indicating similar resolution values across NPA and NHA-based illumination schemes.

**Supplementary Note 2: Control Comparisons Evaluating Reconstruction Deficiencies in Glass and Planar Gold**

In the absence of structured near-field illumination, as is the case for both glass and planar gold substrates, the acquired sub-images lack the spatial modulation required to encode high-frequency information beyond the diffraction limit. As shown in Figure S5Ai–Ci, the diffraction-limited images acquired on glass, planar gold, and nanohole-array substrates all remain constrained by the microscope's resolution limit. However, the spatial distribution and background signal differ between substrates. Following Blind-SIM reconstruction (Figure S5Aii–Cii), the differences become more pronounced.

For the glass substrate (Figure S5Aii), the reconstruction contains diffuse, non-physical high-frequency features and pronounced artefacts that do not correspond to discrete emitters in the diffraction-limited image. A similar behaviour is observed for the planar gold substrate (Figure S5Bii), where the reconstruction produces non-physical high-frequency detail that is not encoded through illumination-induced spatial modulation. Instead, it arises from the reconstruction process itself, despite the absence of any periodic nanostructure capable of generating structured near-field illumination.

In contrast, the nanohole array reconstruction (Figure S5Cii) yields spatially localised and physically meaningful emitter features across the field of view, consistent with true structured illumination generated by the periodic plasmonic nanohole architecture. This provides strong evidence that the observed resolution enhancement is physically encoded by the nanostructured substrate, rather than arising from reconstruction artefacts.

The Blind SIM reconstruction algorithm still converges to a solution in the absence of structured illumination[3]. This behaviour arises from the formulation of Blind SIM as a bilinear inverse problem, in which both the fluorophore distribution $\rho(\mathrm{r})$and the illumination patterns $I_l(\mathrm{r})$are jointly estimated from the measured intensities $M_l(\mathrm{r})$. The forward imaging model is given by:

$$M_l(\mathrm{r}) = \int_\Omega \rho(\mathrm{r}')\, I_l(\mathrm{r}')\, h(\mathrm{r}-\mathrm{r}')\, d\mathrm{r}'$$

where $h$represents the detection point spread function (PSF), and $\Omega$denotes the object domain.

The reconstruction is obtained by minimising the residual:

$$F(\rho, \{I_l\}) = \sum_l \parallel M_l - A(\rho \cdot I_l) \parallel^2$$

To stabilise this highly underdetermined problem, positivity constraints are enforced by reparameterising both the fluorophore density and illumination patterns as squares of real-valued functions:

$$\rho(\mathrm{r}) = \xi^2(\mathrm{r}), I_l(\mathrm{r}) = i_l^2(\mathrm{r})$$

This ensures that both quantities remain non-negative during optimisation. However, as demonstrated by Mudry et al., enforcing positivity can lead to the artificial recovery of high-frequency features that are not present in the measured data. In the absence of structured illumination, these constraints can produce visually plausible but non-physical super-resolved features. This explains why Blind SIM reconstructions obtained on glass and planar gold substrates may appear to contain features beyond the diffraction limit, despite the absence of true resolution enhancement. These results highlight the importance of distinguishing algorithmic artefacts from physically encoded spatial frequency information.

While planar plasmonic films have been experimentally demonstrated to support surface plasmon excitation, this only occurs when appropriate coupling conditions are met to overcome the momentum mismatch, typically through the use of prism- or grating-based configurations [4]. In the absence of such coupling mechanisms in the present system, no plasmon modes are excited, and therefore no structured near-field illumination is generated. In contrast, the engineered nanohole array developed in this work provides intrinsic momentum matching through its periodic geometry, enabling direct excitation of both propagating and localised plasmonic modes. This results in strong confinement and nanoscale spatial modulation of the electromagnetic field, generating the high-spatial-frequency intensity variations required for SIM reconstruction without the need for additional coupling architectures.

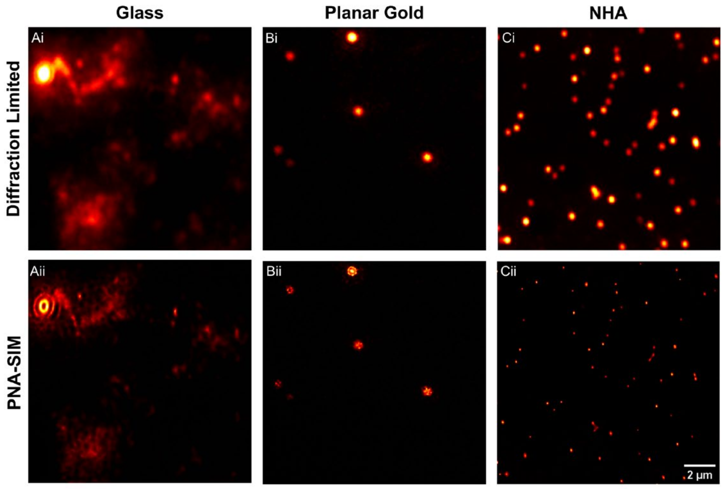


**Figure S5: Super-resolution reconstruction across different substrates.** Comparison of diffraction-limited and reconstructed super-resolution images of 100 nm fluorescent polystyrene nanoparticles on **A** Glass **B** Planar Gold, and **C** NHA substrates. The top row (**Ai–Ci**) shows diffraction-limited images, while the bottom row **Aii–Cii** presents their corresponding super-resolution reconstructions. Unlike Glass and Planar Gold, which fail to provide effective super-resolution reconstruction, the NHA substrate exhibits significant enhancement, demonstrating the critical role of nanostructured confinement in achieving high-resolution imaging.

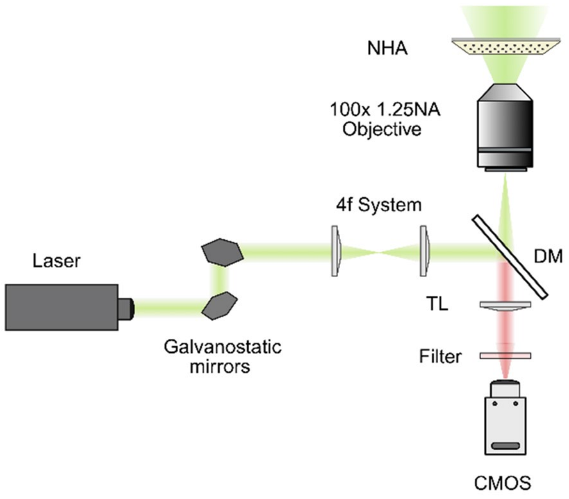


**Figure S6: Simplified schematic of the experimental imaging setup.** A pair of galvanostatic mirrors steer a p-polarised laser beam (λ = 561 nm) through a 4f optical relay system. The beam is reflected by a dichroic mirror (DM) and focused onto the nanohole array (NHA) sample through a high-numerical-aperture (100×, 1.25 NA) objective lens. Fluorescence emission is collected back through the same objective and passes through the dichroic mirror, tube lens (TL), and a fluorescence filter before detection on a CMOS camera. The setup is constructed from modular optomechanical components, enabling customizable structured near-field illumination through angular scanning of the back focal plane.

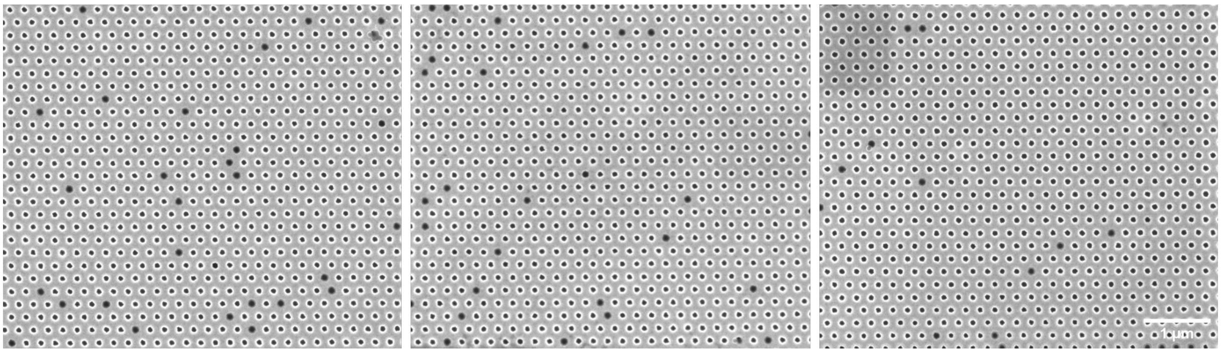

**Figure S7: SEM validation of polystyrene nanoparticle registration within nanoholes.** Across three fields of view, SEM imaging confirms nanoparticle confinement within discrete nanoholes.

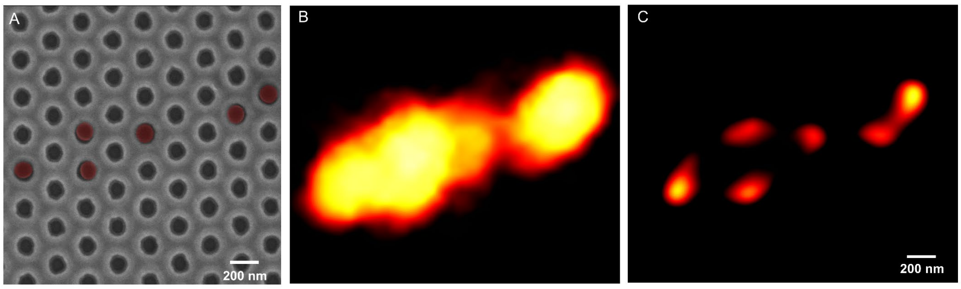


**Figure S8: Correlative validation of nanoparticle localisation and NHA-SIM reconstruction. A** Scanning electron microscopy (SEM) image of the plasmonic nanohole array and fluorescent polystyrene nanoparticles, showing preferential localisation of particles within and around nanohole apertures. False colour overlay to enhance visibility of individual nanoparticles and highlight confinement at nanohole sites. **B** Diffraction-limited fluorescence image of the same region, in which closely spaced particles are not spatially resolved. C Reconstructed NHA-SIM image of the corresponding field of view, resolving discrete emitters consistent with individual nanoparticles.

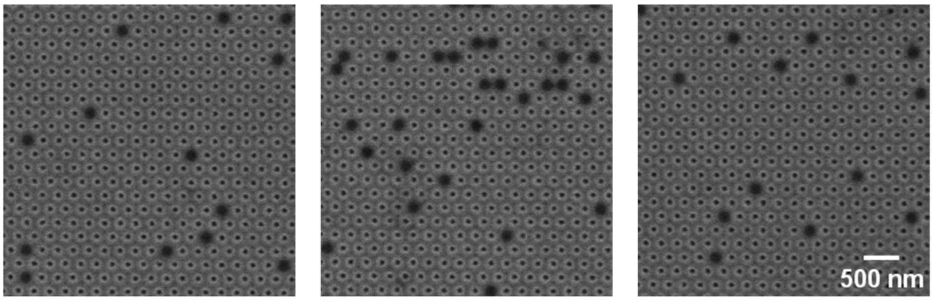


**Figure S9: Nanohole localisation and registration of 200 nm fluorescent particles.** Representative scanning electron microscopy (SEM) images of larger polystyrene beads (200 nm nominal diameter) deposited under identical drop casting conditions. Despite the increased particle size, particles continue to exhibit preferential localisation relative to the nanohole geometry, indicating that the confinement behaviour is robust across different size regimes.

**Table S1**: particles and antibodies information used in this study.

| **Particle/ Antibody** | **Vendor** | **Cat No.** | **Lot No.** | **Labelling** | **Notes** |
|---|---|---|---|---|---|
| Dil label PC liposomes | CD Biosciences | CAT:02600IF-DI | CDJ705 | DiL Membrane Dye | - |
| Lyophilised mCherry Fluorescent EVs from HEK293 cells | Hansa Biomed | HBM-HEK-mCHERRY-9-EGFP-63 | 280425 | CD9 (mCHERRY) & CD63 (EGFP) | - |
| Lyophilised sEVs from HEK293 CL | Hansa Biomed | HBM-HEK293-100/2 | 211123 | Unlabelled | - |
| Lyophilised sEVs from HT29 | Hansa Biomed | HBM-HT29-100/2 | 120423 | Unlabelled | - |

| Particle/ Antibody | Vendor | Cat No. | Lot No. | Labelling | Notes |
|---|---|---|---|---|---|
| EpCAM | Abcam | ab237395 | 1038200-11 | Alexa Fluor 594 | Clone: EPR20532-225<br>Source: Rabbit/ IgG |

**Abbreviations:** DiI, 1,1′-dioctadecyl-3,3,3′,3′-tetramethylindocarbocyanine perchlorate; PC, phosphatidylcholine; EVs, extracellular vesicles; mCherry, monomeric red fluorescent protein Cherry; HEK293, human embryonic kidney 293 cells; HT29, human colorectal adenocarcinoma cells; EpCAM, epithelial cell adhesion molecule.

**Supplementary Note 3: Nanoparticle Tracking Analysis (NTA) of Liposomes and Extracellular Vesicles (EVs)**

The particle concentration and size distribution of DiL liposomes and EpCAM labelled EVs were measured by ZetaView PMX-420 QUATT instrument (Particle Metrix GmbH, Ammersee, Bavaria, Germany) in scatter modes with software version 8.05.16_SP3. All measurements were conducted according to the manufacturer's recommendations. The instrument was first calibrated with the 100 nm PS standard beads. Throughout the measurements, the instrument's temperature control was set to 21.5°C. Before sample injection, the flow cell was flushed with fresh Milli-Q water and then primed with fresh PBS to avoid turbulent drift. Samples were diluted with particle-free 1× PBS to reach the optimal particle concentration within the manufacturer-recommended measurement range (approximately $5 \times 10^6$–$1 \times 10^8$ particles/mL). Particle number and size distribution were counted at 11 individual positions inside the measuring cell under a sensitivity of 80 and a shutter value of 100 (Figure S9A). Other analysis parameters were set as follows: max area 1000, min area 10, min brightness 30, nm/class: 10.

For Lyophilised dye-labelled Fluorescent EVs analysis, fluorescence NTA measurements were performed using a ZetaView Evolution QUATT (equipped with four lasers: 405, 488, 520, and 640 nm) and corresponding long-pass filters with cutoff wavelengths of 410, 500, 550, and 660 nm. No instrument calibration was required prior to measurement for this system. Samples were diluted with particle-free 1× PBS to achieve concentrations within the manufacturer-recommended measurement range (approximately $10^5$ -$10^9$ particles/mL). A 520-nm laser illuminated the particles, and the scattered light was collected. Particle number and size distribution were counted using concentration-scanning technology with a fluorescence sensitivity of 95, a scatter sensitivity of 40, and a shutter value of 100 (Figure S9B). Other analysis parameters were set as follows: frame rate 62.5, recording length 720, min trace length 3, minimum area 0.5, maximum area 500. The specifications of both models are summarised in Table S2. A summary of the results is provided in Table S3.

**Table S2**. Overview of device specifications.

| Device | PMX-420 QUATT | Evolution QUATT |
|---|---|---|
| Camera | High-sensitivity CMOS with 1920 × 1080 pixels | High-sensitivity CMOS with 1280 × 960 pixels |
| Laser | 405, 488, 520, and 640 nm | 405, 488, 520, and 640 nm |
| Flow Cell | 11 individual positions | Concentration Scanning |

| | | |
|---|---|---|
| Calibration | 100 nm PS standard beads | NA |
| ZetaView Software | 8.05.16_SP3 | 1.0.4.6 |

**Table S3:** Particle size and concentration measured by NTA

| Sample Name | Measurement Mode | Median Size (nm) | Concentration in stock solution (particle/mL) | Buffer used in NTA and NHA-SIM |
|---|---|---|---|---|
| PS nanoparticle standard | Scatter | 106.2 | $3.3 \times 10^{13}$ | MilliQ water |
| DiL Liposome | Scatter | 108 | $1.01 \times 10^{13}$ | 0.22 µm filtered 1×PBS |
| Lyophilised mCherry Fluorescent EVs | 550 nm | 120.4 | $3.32 \times 10^{10}$ | 0.22 µm filtered 1×PBS |
| Lyophilised sEV Standards HEK293 | Scatter | 120.6 | $1.30 \times 10^{11}$ | 0.22 µm filtered 1×PBS |
| Lyophilised sEVs Standards HT29 | Scatter | 137.8 | $4.10 \times 10^{11}$ | 0.22 µm filtered 1×PBS |
| EpCAM-labelled HEK293 EVs | Scatter | 148.4 | $5.00 \times 10^{8}$ | 0.22 µm filtered 1×PBS |
| EpCAM-labelled HT29 EVs | Scatter | 132.3 | $6.00 \times 10^{8}$ | 0.22 µm filtered 1×PBS |

**Abbreviations:** DiI, 1,1′-dioctadecyl-3,3,3′,3′-tetramethylindocarbocyanine perchlorate; EVs, extracellular vesicles; HEK293, human embryonic kidney 293 cells; HT29, human colorectal adenocarcinoma cells; EpCAM, epithelial cell adhesion molecule.

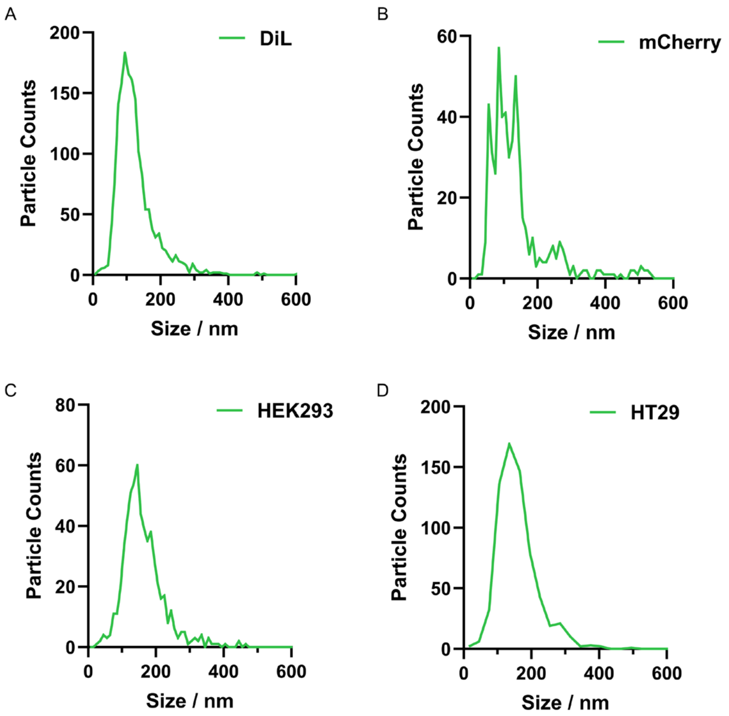


**Figure S10: Nanoparticle tracking analysis (NTA) of liposome and extracellular vesicle (EV) samples used throughout this study. Size distributions obtained from NTA are shown for: A** DiL-labelled liposomes measured in scatter mode. **B** mCherry-labelled EVs measured in fluorescence mode. **C** EpCAM-labelled HEK293-derived EVs measured in scatter mode. **D** EpCAM-labelled HT29-derived EVs measured in scatter mode. Particle counts are plotted as a function of hydrodynamic particle diameter.

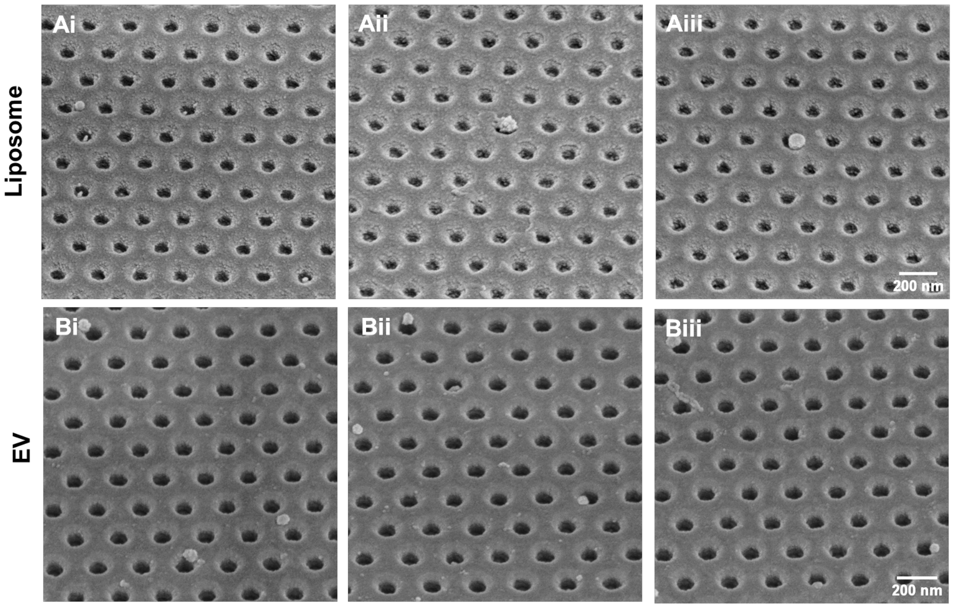


**Figure S11: SEM validation of single-particle occupancy in nanohole arrays. Ai–Aiii.** Scanning electron microscopy (SEM) images of DiI-labelled liposomes confined within the plasmonic nanohole array using the capillary-assisted assembly mechanism, showing representative fields of view with predominantly single-particle occupancy per nanohole. **Bi–Biii**. Corresponding SEM images of mCherry-labelled EVs under identical conditions, demonstrating consistent particle localisation and confinement across multiple regions of the array.

**Supplementary Note 4: SEM Workflow and Sample Preparation Methodology**

SEM was used to characterise the morphology and surface features of the fabricated nanohole arrays and to visualise extracellular vesicles confined within them. Samples were mounted on aluminium stubs using conductive carbon tape or copper tape. Imaging was performed on a NanoSEM 230 field-emission scanning electron microscope (FE-SEM) operated at an accelerating voltage of 5 kV. Liposome and EV samples imaged were prepared using a standard fixation protocol.

Liposome and EV samples were prepared using a standard fixation and dehydration protocol. Briefly, samples were fixed with paraformaldehyde and/or glutaraldehyde in phosphate-buffered saline (PBS), then rinsed in PBS. Post-fixation with osmium tetroxide was performed to enhance membrane contrast. Samples were then dehydrated through a graded ethanol series (30–100%) and dried using hexamethyldisilazane (HMDS). Before imaging, the samples were coated with a 10nm layer of platinum to reduce charging effects.

**Supplementary Note 5: Benchmarking NHA-SIM Resolution Against Diffraction-Limited and Commercial TIRF Microscopy**

To verify that the observed resolution enhancement is intrinsic to NHA-SIM, diffraction-limited widefield (WF) and reconstructed images of mCherry-labelled EVs  were benchmarked against a commercial TIRF microscopy platform (Nikon super-resolution microscope N-STORM) using open-source decorrelation analysis[2]. Representative images (Figure S11Ai–Ci) and corresponding decorrelation analysis (Figure S11Aii–Cii) are shown. mCherry EVs were adsorbed onto Poly-l-lysine solution functionalised substrates, with NHA-SIM imaging performed on gold nanohole array (NHA) chips and TIRF imaging conducted on standard No. 1.5 glass coverslips. All measurements were performed at a concentration of $1 \times 10^9$ particles $mL^{-1}$.

WF NHA-SIM images exhibited a resolution of 462.43 ± 33.75 nm. Following reconstruction, NHA-SIM achieved 130.64 ± 0.30 nm, indicating enhanced spatial frequency support. The commercial TIRF system yielded a value of 387.47 ± 14.68 nm. While improved performance relative to the WF system is expected for TIRF due to its evanescent-field excitation and reduced background, the superior resolution achieved with NHA-SIM demonstrates that the enhancement arises from structured near-field illumination and reconstruction, rather than system-specific effects.

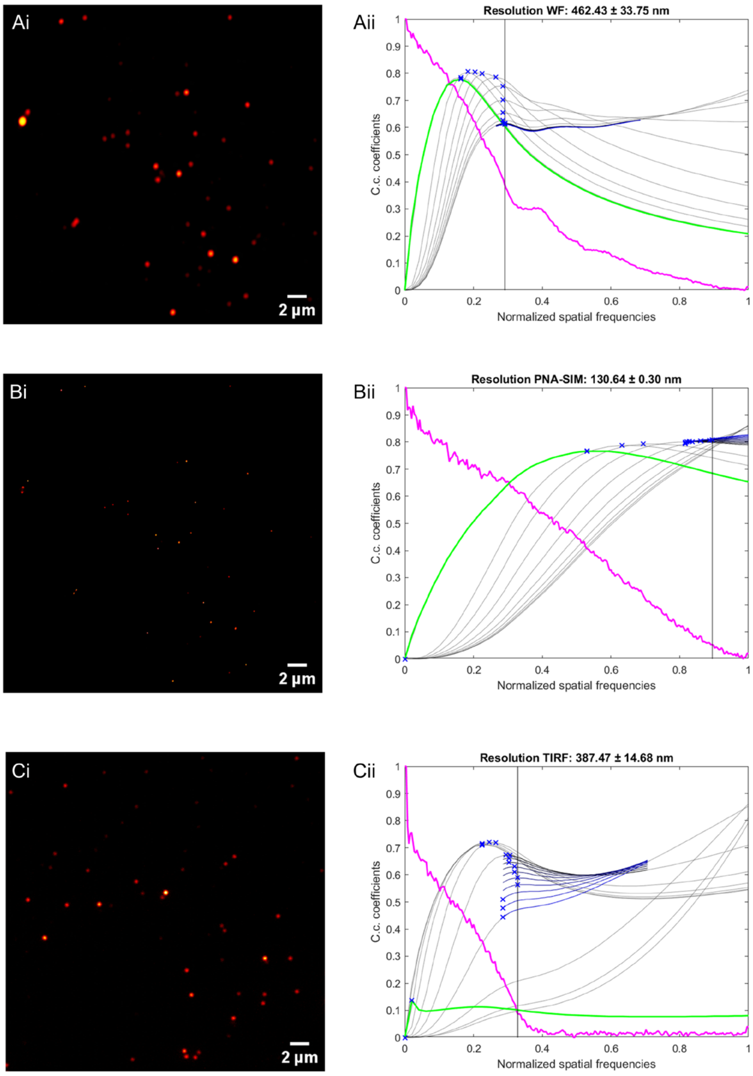


**Figure S12: Resolution benchmarking of NHA-SIM against commercial TIRF. Ai–Ci** Representative fluorescence images of mCherry-labelled EVs acquired using WF NHA-SIM (**Ai**), reconstructed NHA-SIM **Bi**, and commercial N-STORM TIRF microscopy **Ci**. (**Aii–Cii**) Corresponding decorrelation analysis curves used to estimate the spatial resolution.

**Table S4:** Limit of Detection (LOD) Comparison with other Single-EV Counting Technologies

| EV Source | Detection technique | Limit of detection (LOD) | References |
|---|---|---|---|
| BT-474 (HER-2 positive) and MDA-MB-231 | Electrochemical detection | $4.7 \times 10^5$ particles/µL | [5] |
| HEK | Interferometric Reflectance Imaging Sensor (SP IRIS) | $3.94 \times 10^6$ and $5.07 \times 10^6$ particles/µL CD81 and CD63 respectively | [6] |
| HT29 EV and HEK293T | Diffraction-limited microscopy - TIRF | $1.8 \times 10^3$ EVs/µL | [7] |

| EV Source | Detection technique | Limit of detection (LOD) | References |
|---|---|---|---|
| Panc-1 | Surface-Enhanced Raman Spectroscopy Nanotags | $2.3 \times 10^3$ particles/µL | [8] |
| HCT116 | Single-molecule confocal (smConfocal) microscopy | $5.7 \times 10^2$ particles/µL | [9] |
| L-02 and PaNC1 | Dynamic Immunoassay for Single tEV surface protein Profiling (DISEP) | $4.6 \times 10^3$ per µL (L-02 cell derived EVs)<br>$6 \times 10^3$ EVs/µL (Plasma-spiked PaNC1 derived EVs) | [10] |
| **HEK** | **NHA-SIM** | **$1.43 \times 10^2$ EVs/µL** | **This work** |

**Abbreviations:** HER2, human epidermal growth factor receptor 2; HEK, human embryonic kidney cells; HEK293T, human embryonic kidney 293T cells; HT29, human colorectal adenocarcinoma cells; HCT116, human colorectal carcinoma cells; BT-474, human breast cancer cell line; MDA-MB-231, human triple-negative breast cancer cell line; Panc-1 (PaNC1), human pancreatic carcinoma cell line; L-02, human cell line; EVs, extracellular vesicles; TIRF, total internal reflection fluorescence; DISEP, Dynamic Immunoassay for Single tEV surface protein Profiling; SP-IRIS, single-particle interferometric reflectance imaging sensor; NHA-SIM, nanohole array structured illumination microscopy.

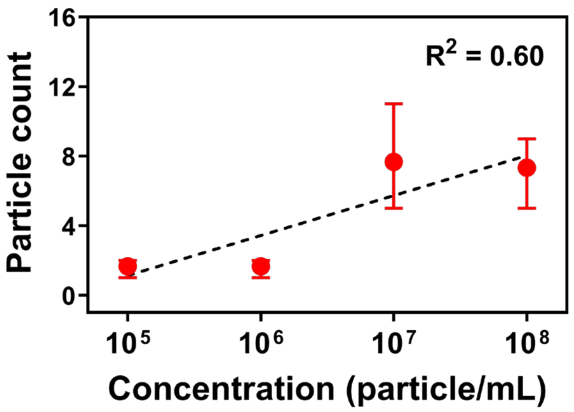


**Fig. S13 Diffraction-limited EV enumeration.** EV counts extracted from diffraction-limited images showed an overall concentration-dependent increase; however, the response exhibited poorer linearity ($R^2 = 0.60$) and deviations from the expected monotonic concentration-dependent trend. This reduced counting fidelity is consistent with nanoscale EV compartmentalisation within the NHA, where closely spaced particles can be detected as merged diffraction-limited features rather than resolved as individual EVs.

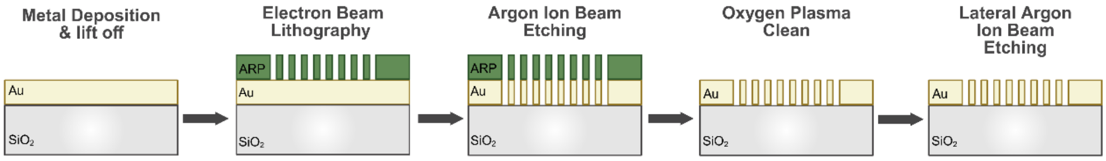


**Figure S14 Fabrication workflow of plasmonic nanohole arrays.** Schematic illustration of the fabrication process flow. A gold (Au) film was first deposited on a $SiO_2$ substrate via Physical Vapour Deposition (PVD) and lift-off, followed by patterning with electron beam lithography (EBL). Argon ion beam etching was subsequently employed to transfer the pattern into the Au layer, with an oxygen plasma treatment used to remove residual resist. A final lateral argon-ion-beam etch was performed to remove redeposited material.
